# Multi-omics characterization of highly enriched human plasma extracellular vesicles

**DOI:** 10.1101/2024.03.25.586695

**Authors:** Huaqi Su, Christopher Fowler, Colin L Masters, Kevin J. Barnham, Gavin E. Reid, Laura J. Vella

**Affiliations:** The Florey Institute, The University of Melbourne, Parkville, Victoria, Australia; School of Chemistry, The University of Melbourne, Parkville, Victoria, Australia; Department of Biochemistry and Pharmacology, The University of Melbourne, Parkville, Victoria, Australia; Bio21 Molecular Science and Biotechnology Institute, The University of Melbourne, Parkville, Victoria, Australia; Department of Surgery, The Royal Melbourne Hospital, The University of Melbourne, Parkville, Victoria, Australia

**Keywords:** Blood, plasma, extracellular vesicles, exosomes, lipidomics, proteomics, proteins, reference, cholesterol esters, mitochondrial cargo, endo lysosomal cargo

## Abstract

Extracellular vesicles (EVs) in blood plasma offer a valuable reservoir of intracellular cellular cargo, making them a promising source of liquid based biomarkers. The molecular cargo of small EVs (sEVs) is of particular interest because some EV subtypes encapsulate cargo from organelles including mitochondria, endosomes, and the autophagy pathways, which are implicated in multiple diseases. However, the complexity of plasma, with its abundance of non-EV particles and plasma proteins, presents challenges for their molecular characterization using mass spectrometry based ‘omics technologies.

Here, we optimised a rigorous method to isolate sEVs from human plasma based on both density and size. Following this, we analysed the protein and lipid content of sEVs from multiple individuals. We demonstrate the advantage of obtaining highly enriched sEVs from plasma for enhancing the detection of protein networks associated with mitochondria and the endosomal network, and also tissue types including the central nervous system. Some of the EV associated proteins reported here have not been detected in plasma, nor plasma sEVs, previously. We show that sphingomyelin lipids are the most abundant lipids in plasma sEVs (33.7 mol% total lipids) and provide the first report on cholesterol ester content. We demonstrate a 16-fold decrease in cholesterol ester lipids in sEVs compared to platelet free plasma and suggest that cholesterol ester content could serve as a valuable measure for assessing the effectiveness of plasma separation protocols or kits in enriching for sEVs.

Our study highlights the benefit of reducing co-isolates from plasma sEV preparations to enable the detection of proteins and lipids with potential biomarker utility, and underscores the need for ongoing development of improved high throughput sEV isolation technologies.

## Introduction

Extracellular vesicles (EVs) are lipid membrane vesicles that are released by all cell types. Depending on their size, cellular origin and molecular constituents, EVs can be classified into a number of categories including microvesicles, apoptotic bodies, exosomes, oncosomes, phagosomes, and even migrasomes [1–4]. EVs circulate in the extracellular environment, are present in bio-fluids such as blood, urine and breast milk [5, 6] and carry a complex cargo including nucleic acids, proteins, lipids, and other metabolites that can report their cell-/tissue-type origin and inform on the pathophysiological conditions of their parental origin. In recent years, there has been increasing interest in examining small EVs (sEVs) derived from endosomes, in blood, with the aim of using sEVs as a source of biomarkers for disease diagnosis, prognosis, and therapeutic monitoring [7–12]. Endosome derived EVs offer an opportunity to interrogate pathways associated with endosome, autophagy, lysosome and mitochondria, outside the cell [13–18]. Investigating blood EVs has the potential to provide valuable insights and serve as an indicator for impairments in these fundamental cellular pathways, which are characteristic of many diseases. However, the complexity of blood and associated difficulties in effectively separating sEVs from the other more abundant blood components (i.e., albumin and lipoprotein particles), have contributed to uncertainty regarding the protein and lipid profiles of sEVs in blood plasma under normal physiological conditions [19]. Chylomicron and low-density lipoprotein particles (LDLs) share a similar size to sEVs, while sEVs have similar density to high-density lipoprotein particles (HDLs) [19, 20]. Due to the overlap in these biophysical features and the significantly higher abundance of lipoprotein particles, sEV isolation via a single conventional method such as ultracentrifugation, density fractionation or size exclusion chromatography (SEC), typically results in co-isolation of plasma lipoprotein particles and soluble proteins [21–23]. Given that lipoproteins are rich in lipids, their co-isolation significantly interferes with molecular profiling of sEVs, leading to diminished lipid and protein coverage and the potential generation of misleading data [24–26]. As a result, the scientific community currently lacks a comprehensive blueprint that outlines both the proteome and lipidome of highly enriched sEVs from plasma. Here we optimized an orthogonal method [27], using density gradient ultracentrifugation (DGUC) to remove LDLs and compared SEC columns for their ability to separate highly abundant proteins and HDLs from DGUC sEVs. By employing this method, we successfully removed abundant plasma proteins, HDLs and LDLs. This resulted in enrichment of plasma sEVs, enabling the identification of more accurate protein and lipid compositions of plasma sEVs. The comprehensive multi-omics data we present serves as a valuable reference for the research community, offering a solid foundation for future analytical studies. Additionally, it serves as a catalyst for the industry to develop high-throughput research kits and biotechnology capable of efficiently removing co-isolates from plasma sEV preparations.

## Materials and Methods

### Blood sample collection, processing, and ethics

Peripheral blood was collected with informed consent from healthy adult donors after overnight-fasting. Blood was drawn through a Safety-Lok, 21G needle collection set (BD Vacutainer) and collected in K2E EDTA tubes (BD Vacutainer). The whole blood collected in K2E EDTA lavender top BD vacutainer was inverted 10 times to mix with EDTA. Within two hrs, blood was processed to separate platelet free plasma (PFP) for plasma sEV isolation. Blood samples were centrifuged at 1,500 x *g* at 20°C for 20 min. Plasma was aspirated and transferred to 15 mL screw cap tubes. Plasma was centrifuged at 2,500 x *g* at 20°C for 10 min, twice, to deplete platelets. The PFP was then aliquoted and stored at −80°C until use. Blood handling and experimental procedures were approved by The University of Melbourne human ethics committee and in accordance with the National Health and Medical Research Council guidelines.

### Plasma small extracellular vesicles (sEVs) isolation via OptiPrep™ density gradient ultracentrifugation (DGUC) and size exclusion chromatography (SEC)

Plasma sEV isolation followed the previously published protocol from Karimi *et al.* [27] with modifications. For SEC column comparison, a pooled PFP sample was used. A schematic of the workflow is shown in Figure 1. Frozen PFP was thawed at 37°C. Before proceeding to plasma EV isolation, a 2 µL aliquot of pooled PFP was retained for protein quantitation and immunoblotting and a 20 µL aliquot was retained for ‘omic characterization.

**Figure 1.**
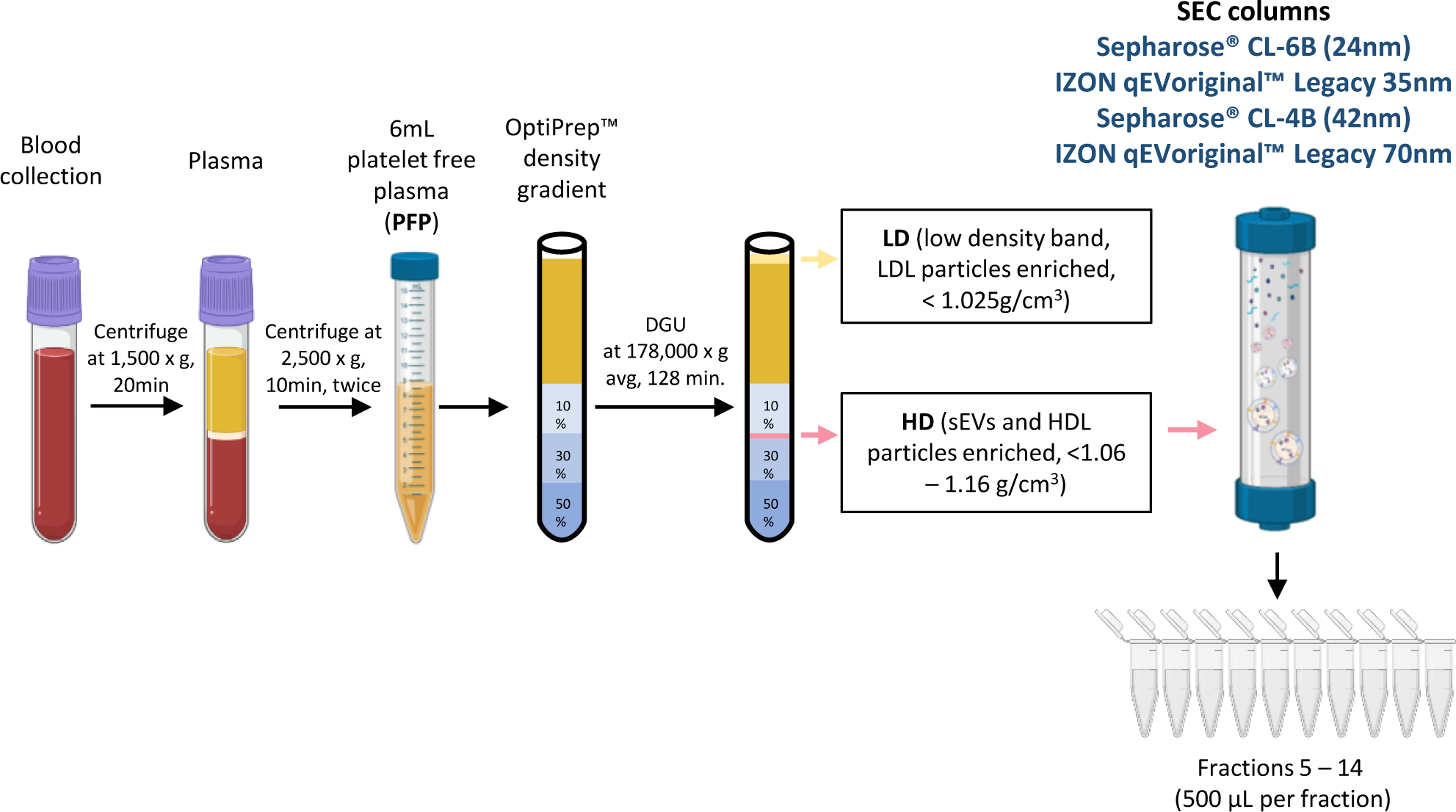
Schematic overview of the plasma sEV isolation workflow: Size exclusion chromatography (SEC) columns were tested for their ability to isolate plasma sEVs and deplete lipoprotein particle following density gradient ultracentrifugation (DGUC). Blood samples were collected in K2E EDTA tubes and spun at 1,500 x g for 20 minutes to separate plasma. Plasma was spun at 2,500 x g for 10 minutes twice to deplete platelets from the plasma. Platelet free plasma (PFP) was collected and stored at −80℃ until use. For sEVs isolation, 6mL of PFP was layered on top of an OptiPrep™ cushion and subjected to 178,000 x g avg, 4℃, for 128 min in a SW40 rotor. HD fractions were pooled before SEC column separation.

An OptiPrep™ density cushion was prepared whereby 2 mL of 50% OptiPrep™ (Sigma Aldrich) phosphate buffer solution (PBS, Thermo Scientific) solution was layered at the bottom of SW40Ti ultra-clear tube (Beckman Coulter), followed by 2 mL of 30% OptiPrep™-PBS solution, and 2 mL of 10% OptiPrep™-PBS solution. Approximately 6.5 mL of PFP was loaded on top of each OptiPrep™ density gradient and a total of 16 density gradients were used. The OptiPrep™ density gradients were subjected to ultracentrifugation at 178,000 x *g* avg, at 4°C for 128 min using a SW 40Ti rotor (Beckman Coulter). After centrifugation, a visible band floating on top of the gradient, containing low density lipoprotein particles (LDLs), was collected, and annotated as low-density band (LD), and a visible band between the 10% and 30% OptiPrep™ layers, containing EVs and high-density lipoprotein particles (HDLs), was collected and annotated as high-density band (HD). All LD and HD collected were pooled respectively, and a 10 µL aliquot from LD and a 20 µL aliquot from HD were retained for protein quantitation and immunoblotting and a 65 µL aliquot from HD was retained for ‘omic characterization. The pooled HD samples were made up to 8 mL with PBS and equally divided into 16 aliquots of 500 µL (4 aliquots for each of the 4 SEC column types) and loaded onto IZON qEVoriginal™ 35nm or 70nm (IZON Science), or home-made Sepharose™ CL-4B or CL-6B (Sigma Aldrich) SEC columns. The home-made Sepharose™ CL-4B and CL-6B SEC columns were packed following a published protocol [28]. Briefly, Sepharose™ CL-4B and CL-6B resins (Sigma Aldrich) were washed with 0.32% citrate in PBS and packed in a 10 mL disposable syringe (Hapool Medical Technology) to a final volume of 10 mL and equilibrated with PBS before use. Separation of content in HD with each SEC column type was repeated four times. For all SEC performance, a fraction of 500 µL was collected, the first 2 mL elution (fractions F1 - F4) was discarded and fractions F5 - F14 were collected for characterization. All columns were washed and equilibrated after elution and before the next aliquot of the HD was loaded.

Fractions from mulitple SEC columns were pooled (fractions F5-F14, a total of 2 mL each) followed by addition of 105.3 µL of 20X PhosSTOP phosphatase inhibitor (Sigma Aldrich) / complete protease inhibitor (including EDTA, Sigma Aldrich) PBS solution. For protein quantitation and immunoblotting, an aliquot of 500 µL from each fraction was mixed with RIPA buffer and sonicated in ice-cold water-bath for 20 min. Protein precipitation was performed by incubating in 20% trichloroacetic acid (TCA) on ice for 30 min followed by centrifugation at 16,000 x *g* for 20 min at 4°C and pellet was washed with 500 µL of –20°C 100% acetone and spun at 16,000 x *g* for 15 min at 4°C, twice, to remove TCA. Protein pellets were air-dried overnight and resuspended in 30 µL PBS containing 2% sodium dodecyl sulphate (SDS), heated at 95°C for 10 min, and proceeded to 20 min water-bath sonication before protein quantitation. The remaining combined SEC fractions were freeze-dried and proceeded to lipid and protein extraction.

### SDS-PAGE and immunoblotting

PFP, LD and HD samples were mixed with phosphatase inhibitor and protease inhibitor PBS solution and RIPA buffer and sonicated in ice-cold water-bath for 20min. Protein content was determined by Pierce™ BCA assay kit (Thermo Scientific) according to the manufacturer’s instructions. PFP, LD, HD samples and equal volume of all SEC protein fractions were prepared in 4X NuPAGE™ LDS Sample Buffer and 10X Bolt ™ sample reducing agent and boiled (10 min, 90°C) followed by centrifugation (16,000 x g, 1 min). Normalized PFP, LD and HD samples (2.3 µg) and equal volume for all SEC fractions were electrophoresed on 4-12% NuPAGE™ Bis-Tris in NuPAGE™ MES SDS Running Buffer (Thermo Scientific) for 60 min at 160 V. Proteins were transferred onto nitrocellulose membranes using an iBlot™ 2 Dry Blotting System (Thermo Fisher Scientific). Total protein profile on the membranes was stained using Revert™ 700 Total Protein Stain following instruction (LI-COR) and imaged on an Odyssey® Fc Imaging System (LI-COR).

After removing staining, membranes were blocked with 1X Blocker™ FL Fluorescent Blocking Buffer (Thermo Scientific) for 1 hr at room temperature followed by 48 hr incubation with primary antibodies (syntenin #ab133267 from Abcam, flotillin-1 #610821 from BD Biosciences; ApoA-1 #3350 from Cell Signaling Technology and ApoB #139401 from Abcam) at 4°C. Membranes were washed 4 times with TBS-T (30 min, room temperature) following a 1 hr, room temperature incubation of the IRDye® 800CW Goat anti-Rabbit IgG secondary antibody (#925-32211, LICOR) or the IRDye® 800CW Goat anti-Mouse IgG secondary antibody (#926-32210, LICOR). All antibodies were diluted in TBS-T. Membranes were washed 4 times with TBS-T (30 min, room temperature). The membranes were visualised on an Odyssey® Fc Imaging System (LI-COR).

### Lpid extraction from plasma and derived fractions

Freeze-dried SEC fraction samples, along with PFP (20 μL) and HD (65 μL) samples, were prepared in 200 µL ice-cold 60% methanol (LCMS grade, EMD Millipore Corporation) containing 0.01% (w/v) butylated hydroxytoluene (BHT, Sigma Aldrich). All samples were sonicated in an ice-cold water bath sonicator for 20 min prior to monophasic lipid extraction following a method previously reported [29, 30]. Briefly, 120 μL of MilliQ water, 420 μL of methanol with 0.01% (w/v) BHT, and a total of 270 μL of chloroform with 0.01% (w/v) BHT (containing internal lipid standards, see below for list of internal standards) were added to all samples, followed by 10 sec vortexing and agitation at 1,000 rpm for 30 min at room temperature. Samples were spun at 14,000 rpm for 15 min at room temperature and supernatants containing lipids were transferred to new tubes. 100 µL of MilliQ water and 400 μL of chloroform:methanol (1:2, v:v) containing 0.01% (w/v) butylated hydroxytoluene (BHT) were added to remaining pellets for re-extraction to maximise lipid recovery following incubation and centrifugation as described above. Lipid extract supernatants from the same extractions were pooled and dried by using a GeneVac miVac sample concentrator (SP Scientific, Warminster, PA, USA). PFP and HD lipid extracts were reconstituted in 400 µL isopropanol:methanol:chloroform (4:2:1, v:v:v, containing 0.01% BHT) while SEC fraction lipid extracts were reconstituted in 200 μL isopropanol:methanol:chloroform (4:2:1, v:v:v, containing 0.01% BHT). The customised isotope labelled internal standard mixture was comprised of the following deuterated lipid standards (Avanti Polar Lipids, Alabaster, AL, USA): 15:0-18:1(d7) PC (204.3 µM), 15:0-18:1(d7) PE (67.6 µM), 15:0-18:1(d7) PS (61.9 µM), 15:0-18:1(d7) PG (37.8 µM), 15:0-18:1(d7) PI (34.1 µM), 15:0-18:1(d7) PA (9.8 µM), 18:1(d7) LPC (45.5 µM), 18:1(d7) LPE (9.9 µM), 18:1(d7) Chol Ester (438.3 µM), 18:1(d7) MG (13.2 µM), 15:0-18:1(d7) DG (8.2 µM), 15:0-18:1(d7)-15:0 TG (59.2 µM), 18:1(d9) SM (52.1 µM), d18:1(d7)-15:0 Cer (27.2 µM).

### Direct infusion nano-electrospray ionization (nESI) - ultrahigh resolution accurate mass spectrometry (UHRAMS) lipidome analysis

10 μL of PFP lipid extract, 20 μL of HD lipid extract and 30 μL of sEVs fraction lipid extracts were aliquoted to individual wells of a twin-tec® 96-well plate (Eppendorf, Hamburg, Germany), dried and then reconstituted in 40 μL (PFP and HD) and 15 μL (sEVs fractions) of isopropanol:methanol:chloroform (4:2:1, v:v:v) containing 20 mM ammonium formate respectively. The 96-well plate was then sealed with Teflon Ultra Thin Sealing Tape prior to mass spectrometry analysis. 10 μL of lipid sample was then aspirated and introduced via nano-ESI to an Orbitrap Fusion Lumos mass spectrometer (Thermo Fisher Scientific, San Jose, CA, USA) using an Advion Triversa Nanomate (Advion, Ithaca, NY, USA) operating with a spray voltage of 1.3 kV and a gas pressure of 0.3 psi in both positive and negative ionization modes. For MS analysis, the RF lens was set at 10%. Full scan mass spectra were acquired at a mass resolving power of 500,000 (at 200 m/z) across a m/z range of 150 – 1600 using quadrupole isolation, with an automatic gain control (AGC) target of 2e5, and a normalized AGC target of 50%. The maximum injection time was set at 100 ms. Spectra were acquired and averaged for 1.5 min. Samples were measured in triplicates.

### Lipid identification, quantitation, and data analysis

‘Sum composition’ level lipid identifications were achieved using LipidSearch software (ver 5.1.6,Mitsui Knowledge Industry, Tokyo, Japan) by automated peak peaking and searching against a user-defined custom database of lipid species (including the deuterated internal standard lipid species). The parent tolerance was set at 3.0 ppm, a parent ion intensity threshold three times that of the experimentally observed instrument noise intensity, and a max isotope number of 1 (i.e., matching based on the monoisotopic ion and the M+1 isotope), a correlation threshold (%) of 0.3 and an isotope threshold (%) of 0.1. The lipid nomenclature used here follows that defined by the LIPID MAPS consortium [31, 32]. Lipid ions detected in at least 2 of 3 technical replicate measurements in any sample were included for quantitation and downstream analysis. Semi-quantitative analysis of the abundances of identified lipid species was performed using an in-house R script, by comparing the identified lipid ion peak areas to the peak areas of the internal standard for each lipid class or subclass (if available, otherwise other structurally similar internal standards). For statistical analysis, missing values were replaced with 1/3 of the minimum value of the entire dataset.

The calculated lipid abundance of individual lipid ions, and the mean summed lipid abundances at the lipid category, class or subclass levels, were normalized to either the total lipid concentration (i.e., mol% total lipid), or total lipid-class concentration (i.e., mol% total lipid class) then used to compare the differences in lipid profiles among PFP, HD and the sEV fractions.

Significant differences in the mean normalized abundances, at the lipid category (mol% total lipid), class (mol% total lipid) and subclass (mol% total lipid class) levels, were determined by ANOVA followed by Sidak’s multiple comparisons test using the GraphPad Prism 9.0 software, with multiplicity adjusted *p* value < 0.05.

### Proteomics sample preparation

After lipid extraction, the remaining protein pellets were air-dried overnight. The PFP, LD and HD pellets were dissolved in 200 µL 5% SDS in PBS solution and SEC fraction pellets were dissolved in 80 µL and heated at 95°C for 10 min with agitation. Protein content was again determined by Pierce™ BCA assay kit (Thermo Scientific) according to the manufacturer’s instructions. Protein reduction, alkylation and digestion was carried out via suspension trapping using the micro S-Trap (Protifi) cartridges following the manufacturer’s instruction with slight modifications [33–35]. Briefly, 3.5 µg protein from each sample was transferred to protein low bind tubes, reduced and alkylated by 10 mM tris (2-carboxymethyl) phosphine (TCEP) and 40 mM chloroacetamide (CAA) in 50mM triethylammonium bicarbonate (TEAB) buffer at 99°C for 5 min. The reaction was then quenched in 1.2% phosphoric acid and a 7X volume of binding buffer (0.1 M TEAB and 90% methanol solution, pH 7.1) was added to each sample to generate the protein particles. Protein suspensions were transferred to micro S-Trap cartridges followed by centrifugation at 4,000 x g for 1 min to remove solvent and trap protein particles. Protein bound to the quartz membrane was washed with 150 µL binding buffer three times and digested with Pierce™ Trypsin Protease MS-Grade (Thermo Scientific) dissolved in 50 mM TEAB buffer with digestion ratio of 1:20, trypsin:protein, w:w. Digestion occurred in a 37°C incubator for overnight. Digested peptides were eluted with three subsequent buffers, 40 µL of 50 mM TEAB buffer, 40 µL 0.2% formic acid and 40 µL 50% aqueous acetonitrile containing 0.2% formic acid. Protein solution was dried and reconstituted in 21.6 µL of 2% acetonitrile containing 0.05% trifluoroacetic acid. PFP, HD, 6B-F10, 35nm-F10, 4B-F10 and 70nm-F11 protein suspensions were processed and analysed in triplicate to assess reproducibility of the workflow.

### Data independent acquisition (DIA)-liquid chromatography tandem mass spectrometry (LC-MS/MS) proteomics analysis

For each sample, 0.7 µg of the peptide digests were introduced to an Acclaim Pepmap nano-trap column (Dionex-C18, 100Å, 75 µm x 2 cm) uisng an isocratic flow rate of 5 µL/min in 2% ACN + 0.05%TFA for 6 min. The trap column was then switched in-line with the Acclaim Pepmap RSLC analytical column (Dionex-C18, 100Å, 75 µm x 50 cm) and peptides then analysed using an Orbitrap Eclipse Tribrid mass spectrometer (Thermo Fisher Scientific). Solvent A comprised 0.1% formic acid (v/v) and 5% dimethyl sulfoxide (DMSO) in water (v/v). Solvent B comprised ACN + 0.1% formic acid (v/v) and 5% DMSO in water (v/v). Each sample was eluted with the following gradient at a flow rate of 0.3 µL/min. The gradient started at 3% to 23% Solvent B over from 6 to 95 min, increased from 23% to 40% Solvent B from 95 to 105 min, increased from 40% to 80% Solvent B from 105 - 110 min, stabilized at 80% Solvent B for 5 min, lowered to 3% Solvent B over 0.1 min and stabilized at 3% until finish at experimental run time of 125 min. Samples were delivered via nano-spray infusion with a spray voltage of 1.9 kV in positive ionisation mode, with the ion transfer tube temperature set ay 275°C. The full scan MS spectra data acquisition parameters included wide quad isolation; desired minimum points across the peak of 6; detector type, orbitrap mass resolution of 120,000; scan range of 350-1400 m/z; maximum injection time, 50 ms; normalized AGC target, 40%; microscans, 1; polarity, positive; and RF lens of 30%. The default charge group state was set as 2. For HCD-MS/MS data acquisition, orbitrap with mass resolution was 30,000; HCD collision energy was 32%, the desired minimum points across the peak was 9, collision energy type, normalized; isolation mode, quadrupole; multiplex ions disabled; window placement optimization disabled; number of scan events, loop count, 20; loop control, 3; loop time, 3; mass range, normal; precursor mass range, 361-1033; scan range, 200-2000 m/z; maximum injection time, 55 ms; normalized AGC target, 2000%; microscans, 1; data type, centroid; polarity, positive; and source fragmentation disabled. Precursor mass range of 360.5 to 1033.5 m/z, with an isolation window of 13.7 m/z, and an overlap window of 1 m/z.

### DIA proteomics data analysis and protein identification

MS raw files were analyzed using the Spectronaut v.14 (Biognosys, Switzerland) with library-free “direct-DIA” approach. The spectra fragment lists were searched against the Swiss-Prot *Homo Sapiens* proteome FASTA library (downloaded on 20220829) for protein identification. The search parameters are as follows. Under Pulsar Search peptide setting, enzymes: Trypsin/P, digest type: specific, max. peptide length 52, min. peptide length: 7, missed cleavages was set as 2. Max variable modifications, 5, fixed modification: carbomidomethyl (C); variable modification: acetyl (protein N-term) and oxidation (M). Under DIA analysis setting, for identification, Q value cut-off was set as 1% for identification at precursor and protein levels. Single hit proteins were excluded, and proteins were defined by stripped sequence. False discovery rates (FDR) for peptide spectrum match (PSM), peptide, and protein group were set to 0.01; Quantification was performed with default settings: identified Qvalue for precursor filtering; imputation strategy and proteotypicity filter were none.

Protein label free quantification (LFQ) method, automatic; quantity MS level, MS2; quantity type, area. Cross-run normalization was enabled, and normalization strategy, automatic. Major protein grouping by protein group ID; minor peptide grouping by stripped sequence; major group quantity by mean peptide quantity; minor group quantity by mean precursor quantity. Default parameters were used for chromatograms (XIC) Extraction, Calibration, Workflow, and Pipeline Mode. After searching and filtering, enrichment analyses for gene ontology, KEGG pathways and Reactome pathway for sEVs enriched fractions were performed using GeneCodis4 with default significance threshold.

## Results

Small EVs and lipoprotein particles have similar physical traits (size and density) so employing methodologies that combine DGUC and SEC is essential for both isolating and enriching plasma sEVs while simultaneously depleting lipoprotein particles and protein co-isolates [19, 20]. To achieve this, we adapted a previously published method (Figure 1) [27]. In brief, platelet free plasma was subject DGUC to remove LDLs from the EVs and HDLs residing in a high-density fraction. Then, to further separate sEVs from HDLs, the high density fraction was applied separately to 4 commonly used SEC columns, the qEVoriginal™ 35nm and 70nm columns, and home-made SEC columns packed with Sepharose™ CL-4B and CL-6B resins (Figure 1) [28]. The eluted fractions were subsequently analysed by immunoblot, and proteomics and lipidomics.

### Immunoblotting results of sEVs isolated using DGUC and SEC columns

Immunoblotting showed that syntenin, an endosome-derived EV marker, was enriched in fractions (F) F8 and F9 from Sepharose™ CL-6B, F7-9 from qEVoriginal™ 35nm, F8 and F9 from Sepharose™ CL-4B, and F7-9 from qEVoriginal™ 70nm, suggesting that these fractions contained sEVs (Figure 2). Consistent with this, another generic EV marker, flotillin-1, was enriched in F7-9 qEVoriginal™ 35nm, and F7-9 qEVoriginal™ 70nm with lower levels in F8 and F9 Sepharose™ CL-6B and F8 and F9 Sepharose™ CL-4B. Importantly, there was negligible ApoB100 detected in the sEV fractions, indicating that LDLs (ApoB100 rich) were efficiently removed by DGUC. The early eluting sEV fractions from the qEVoriginal™ 70nm SEC column were found to be cleaner than the sEV fractions eluted from the other 3 columns. The DGUC qEVoriginal™ 70nm SEC sEV fractions contained less free protein (as seen by the total protein stain) and contained less HDLs (ApoA-I) than the sEVs isolated with the other 3 columns. Therefore, DGUC followed by qEVoriginal™ 70nm SEC was found to be the most effective at separating LDLs, reducing free protein, and removing HDLs from sEVs.

**Figure 2.**
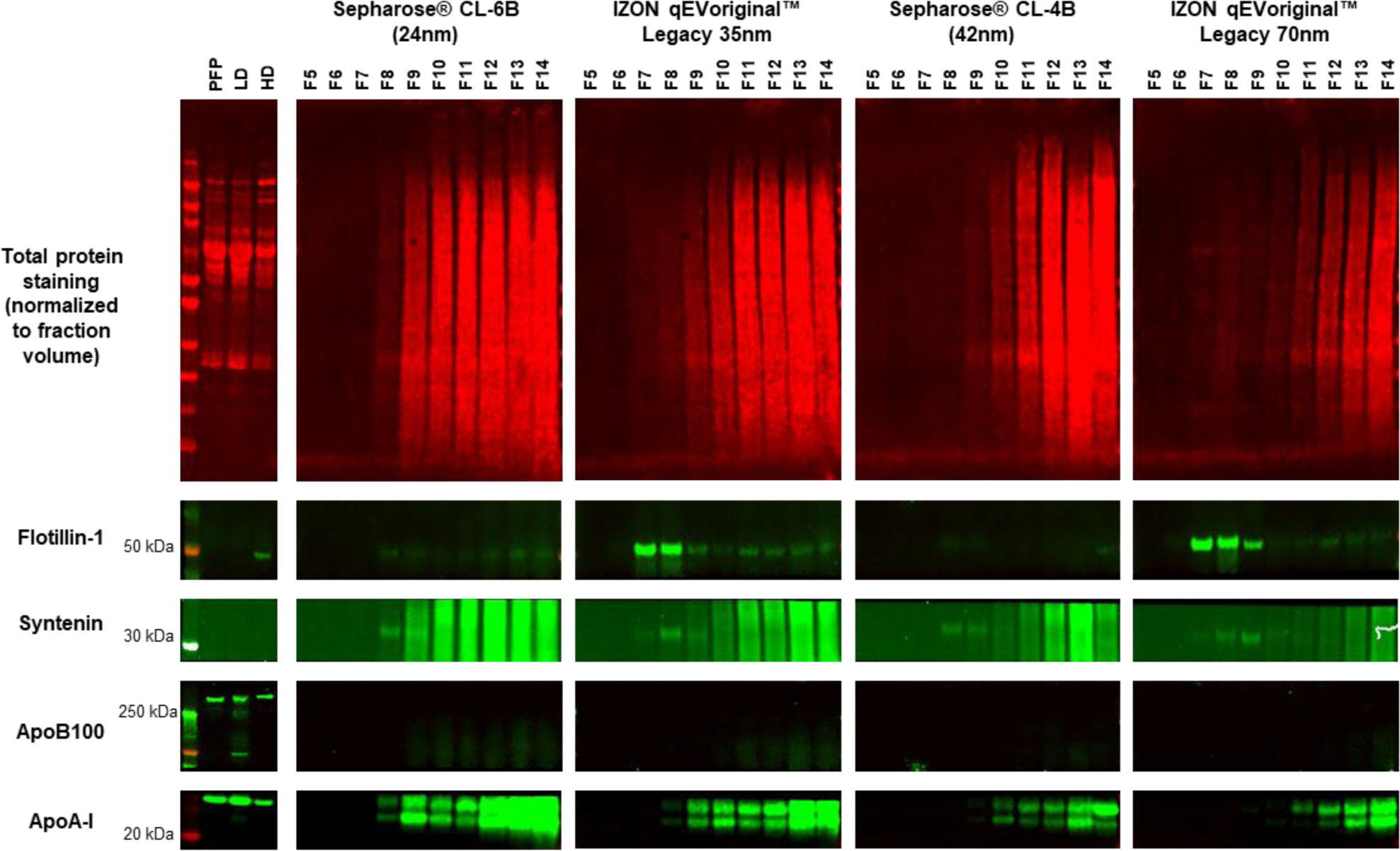
Immunoblot characterization of plasma sEVs isolated by density gradient ultracentrifugation (DGUC) followed by size exclusion chromatography (SEC). PFP, LD and HD samples (2.3 µg) and SEC fractions (equal volume) were electrophoresed for 60 minutes at 160 V. Total proteins were stained and imaged, followed by immunoblotting with sEV markers, flotillin-1 and syntenin, and ApoB100 (LDL marker) and ApoA-I (HDL marker).

### Proteomic characterization of sEVs isolated using DGUC and SEC columns

The presence of contaminating proteins and lipoprotein particles in plasma poses challenge for detection and identification of low abundant sEV associated proteins (present in much lower scale in plasma) by mass spectrometric analysis. To evaluate the benefit of removing abundant plasma proteins and lipoproteins from sEV preparations, the PFP, LD, HD, and fractions collected from DGUC followed by SEC were submitted for DIA proteomic analysis. A total of 490, 429 and 619 proteins were identified in the PFP, LD and HD, respectively. In contrast, proteome analysis of the Sepharose™ CL-6B, qEVoriginal™ 35nm, Sepharose™ CL-4B and qEVoriginal™ 70nm sEVs resulted in the identification of a combined 994, 1694, 1495 and 1736 proteins, respectively (Figure 3A, Supplementary Table 1). This demonstrates that by effectively separating LDLs, reducing free protein, and removing HDLs from sEVs using DGUC followed by qEVoriginal™ 70nm SEC, a greater number of sEV proteins may be detected by MS. Supplemental Figure 1 shows the reproducibility and robustness of the protein preparation and DIA proteomic analysis workflow.

**Figure 3.**
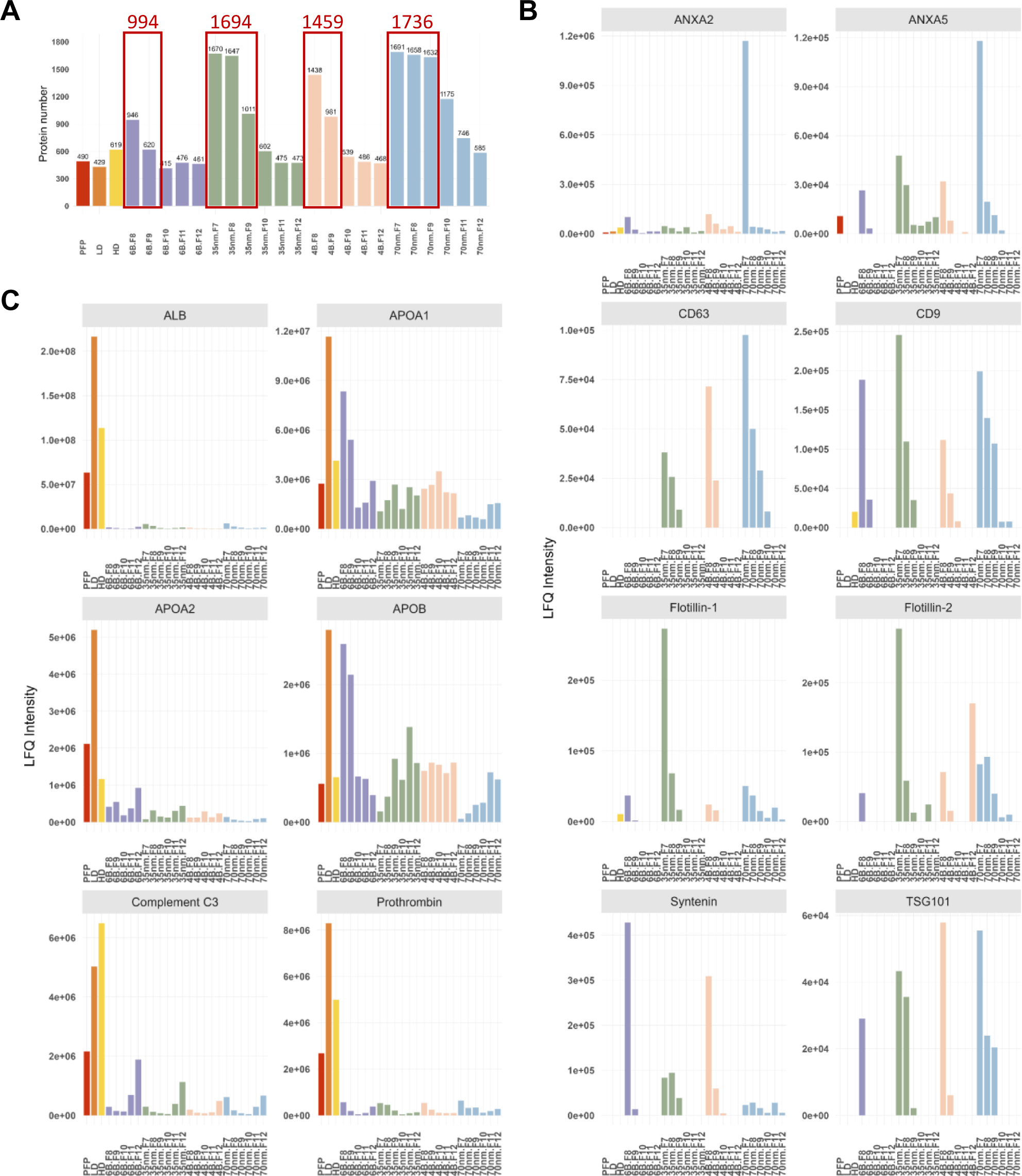
SEC columns vary in their ability to enrich for sEVs and to deplete non-EV particles and abundant plasma proteins following DGUC. **A.** Number of protein identifications in each sample. The sEVs enriched fractions are highlighted by the red boxes and the numbers in red indicate the number of proteins identified in these fractions. **B.** The LFQ comparison of selected EV markers. The sEVs enriched fractions contained higher level of EV markers (ANXA2, ANXA5, CD63, CD9, flotillin-1, flotillin-2, syntenin and TSG101). **C.** The LFQ comparison of contaminating proteins. The sEVs enriched fractions contained substantially low intensity of contaminant proteins (ALB, APOA1, APOA2, APOB, complement C3 and prothrombin). PFP, platelet free plasma, in red; LD, low density band, in orange; HD, high density band, in yellow; 6B = Sepharose™ CL-6B, in purple; 35nm = qEVoriginal™ 35nm, in green; 4B = Sepharose™ CL-4B, in pink; 70nm = qEVoriginal™ 70nm in blue.

The LFQ intensity comparison of some general EV markers and contaminant proteins in the PFP, LD, HD and SEC fractions are shown in Figure 3B and Figure 3C respectively. The sEV enriched fractions contained EV proteins (ANXA2, ANXA5, CD63, CD9, FLOT1, FLOT2, SDCBP and TSG101) with differences seen between the SEC columns (Figure 3B) in regards to the relative abundance of these markers. For example, CD63 was present in the sEVs eluted from all SEC columns, except the Sepharose™ CL-6B column. While the sEVs isolated using Sepharose™ CL-6B column contained minimal CD63, sEVs isolated from the Sepharose™ CL-6B column and Sepharose™ CL-4B column contained higher syntenin relative to sEVs isolated from the two IZON qEVoriginal™ SEC columns. Additionally, sEVs from all SEC columns yielded comparable protein levels of CD9 and TSG101. The LFQ intensity of potential co-isolated contaminant proteins (ALB, APOA1, APOA2, APOB, complement C3 and prothrombin) were lower in the sEVs fractions eluted from qEVoriginal™ 35nm and qEVoriginal™ 70nm (Figure 3C) relative to the other two SEC columns. While the depletion of ALB, APOA2, complement C3 and prothrombin were all observed in sEV fractions from the Sepharose™ CL-6B and Sepharose™ CL-4B columns, the levels of ApoA1 and ApoB proteins were comparatively higher (Figure 3C). Among the SEC columns, the qEVoriginal™ 70nm SEC best enriched sEVs following DGUC, and the sEVs had the highest levels of EV markers and the lowest levels of plasma protein and lipoprotein contaminates. The immunoblot and proteomic data suggest that combining DGUC with the qEVoriginal™ 70nm enabled the isolation of the highest purity plasma sEVs.

Additional analysis of the identified proteins using gene ontology, KEGG pathway, and Reactome enrichment analysis further confirmed the efficient enrichment of sEVs using this approach. Figure 4A shows the top 10 gene ontology enriched biological processes, molecular functions, and cellular components, while Figure 4B highlights the Top 10 KEGG pathways associated with the central nervous system (CNS) and cellular processing (ribosome, proteasome, and phagosome). In the Reactome enrichment analysis, the sEVs proteome was found to be associated with axon guidance, nervous system development, immune system and platelet activity and regulation (Figure 4C).

**Figure 4.**
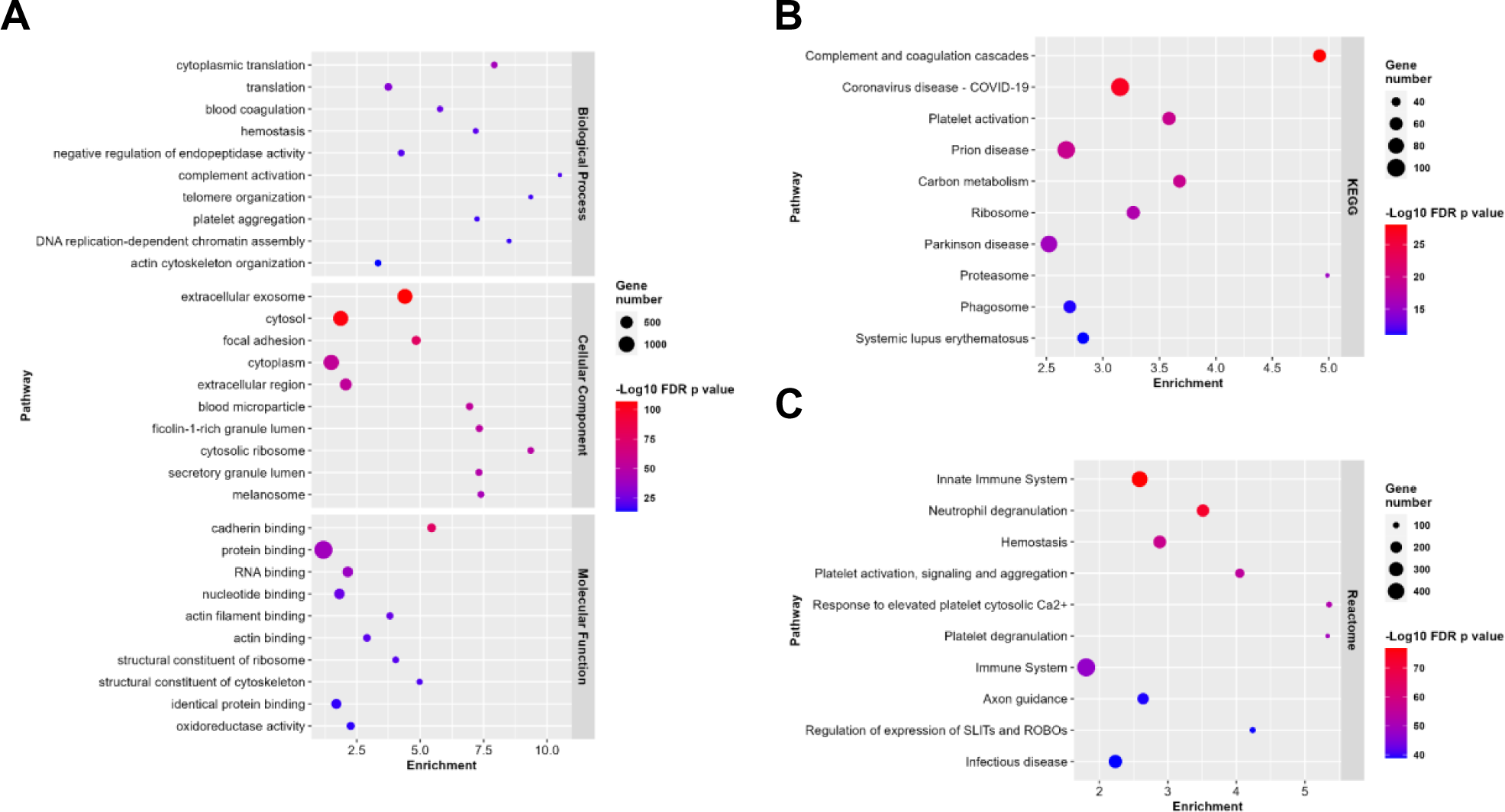
Enrichment analysis of the proteome of highly enriched plasma sEVs. **A.** Gene ontology enrichment analyses including Top 10 biological process, cellular component and molecular function pathways. **B.** Top 10 enriched KEGG pathways. **C.** Top 10 enriched Reactome pathways. The sEVs were isolated from plasma using DGUC and qEVoriginal™ 70nm SEC (Fractions F7-9).

The highly enriched sEV proteome was compared to previously published plasma EV proteomes reported by Karimi *et al.* [27] and EVs isolated using PS-affinity capture by Muraoka *et al.* [36] (Supplementary Figure 2A). These two studies were chosen for comparison as the Karimi *et al.* study is regarded as the most rigorous to date in effectively demonstrating enrichment of EVs and depletion of plasma co-isolates, while the Muraoka *et al.* study presented the highest protein coverage in blood EVs, albeit with caveats around the purity of the EVs [36]. Of the 177 proteins that were uniquely identified in our sEVs (Supplementary Table 2), interactive network analysis showed these proteins are associated with the proteasome accessory complex, ribosome, ribonucleoprotein complex, DNA dependent protein kinases, paraspeckles and spliceosome that involve in regulation of transcription (Supplemental Figure 2B), and mitochondria associated protein clusters (Supplemental Figure 2C), namely, mitochondrial respiratory chain complex, mitochondrial respirasome and oxidoreductase complex.

### Lipidomic characterization of sEVs isolated using DGUC and qEVoriginal™ 70nm SEC column

Lipidomic analysis was also performed on the PFP, HD and individual qEVoriginal™ 70nm sEVs fractions F7, F8 and F9. Lipids were identified from four main lipid categories, including glycerophospholipids (GP), sphingolipids (SP), glycerolipids (GL) and sterol lipids (ST). A complete list of the individual identified lipids and their semi-quantitative abundances can be found in Supplemental Table 3. The number of lipids and the mean summed lipid abundances at the lipid category and lipid class levels, expressed as mol% total lipid and mol% lipid class, are summarized in Supplemental Table 4. The sEV enriched fractions F7, F8 and F9 contained significantly less glycerophosphocholine (PC) than PFP and HD (Figure 5). lycerophosphoethanoamine (PE) lipids were found to be enriched in the sEV fractions F7 (18.35 mol% total lipids) and F8 (23.20 mol% total lipids) compared to PFP, where PE lipids made up 2.53 mol% total lipids. Glycerophosphoserine (PS) lipids were one of the most enriched lipid classes in the sEV fractions, e.g., F7 where they comprised 28.82 mol% of the total lipid abundance. Similar to the PE lipids, sphingomyelin (SM) lipids were enriched in sEVs F7 and F8 compared to PFP and HD, representing 9.98 mol% of the total lipids in F7 and 8.67 mol% of the total lipids inF8) Cholesteryl ester (CE) lipids, major lipid components in lipoprotein particles, were highly abundant in PFP (37.93 mol% total lipids) and HD (14.83 mol% total lipids), but were significantly depleted in all sEVs fractions, i.e., 2.20 mol% total lipids, 3.76% total lipids and 8.09 mol% total lipids in F7, F8 and F9 respectively, indicating the efficient removal of lipoprotein particles from sEVs. This lipidome data shows that highly enriched plasma sEVs are rich in PE, PS and SM lipids, consistent with what has been reported previously for sEVs isolated from cell lines [37, 38] and tissue [30] and are highly depleted of lipoprotein predominating CE lipids, indicative of the high purity of sEVs isolated using this approach.

**Figure 5.**
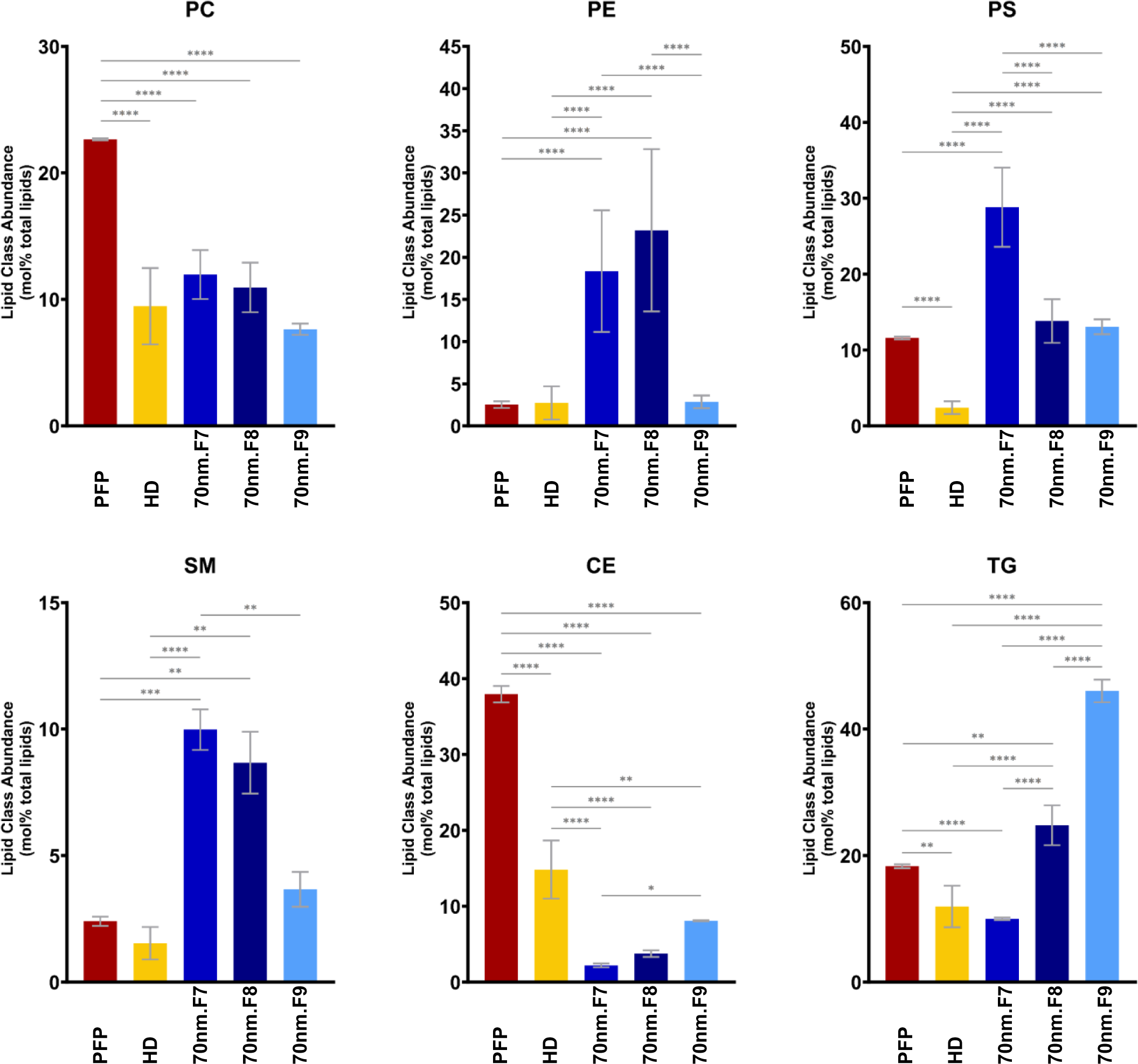
Comparison of mol% total lipid abundance in PC, PE, PS, SM, CE and TG lipid classes in PFP, HD, and highly enriched sEVs. Data represent mol% total lipid abundance mean ± standard deviation. Statistical significance was determined by ANOVA followed by Sidak’s multiple comparison test, with multiplicity adjusted *p* value < 0.05, *p < 0.05; **p < 0.01, ***p < 0.001, ****p < 0.0001. PFP = platelet free plasma, HD = high density band. sEVs were isolated from plasma using DGUC followed qEVoriginal™ 70nm (fractions F7-9). PC = glycerophosphocholine, PE = glycerophosphoethanoamine, PS = glycerophosphoserine, SM = sphingomyelin, CE = cholesteryl ester, TG = triglyceride.

### Proteomic characterization of sEVs isolated from three young and healthy female volunteers using DGUC and qEVoriginal™ 70nm SEC column

With confidence in the methodology to isolate highly enriched sEVs from plasma, we embarked on investigating the protein and lipid composition of these vesicles with the goal of determining if sEVs have a defined and reproducible composition between individuals. We isolated plasma sEVs from three young healthy female volunteers using the optimized method described above, combining DGUC and qEVoriginal™ 70nm SEC fractions F7-F9. On average, 1 mL of PFP yielded sEVs equivalent to 3.71 ± 0.28 µg protein.

Immunoblotting analysis showed that sEVs from the three individuals were enriched in flotillin-1 and syntenin, had limited ApoB100 and contained negligible ApoA protein (Figure 6A). Proteomic analysis of the combined F7, F8 and F9 sEV fraction resulted in the identification of 167 proteins, a count lower than the combined proteome observed from the individual fractions reported above (Figure 6B). This difference may be attributed to the additional proteome complexity introduced by the combination of F7, F8 and F9 fractions, which hindered the dymanic range during mass spectrometric analysis. However, the protein content of the sEVs from these combined fractions was very similar between individuals, as shown by the Venn diagram in Figure 6C, with clear enrichment in EV proteins (Figure 6D) and depletion of blood contaminants (Figure 6E) compared to PFP across individuals, similar to that seen in the individual fractions reported above.

**Figure 6.**
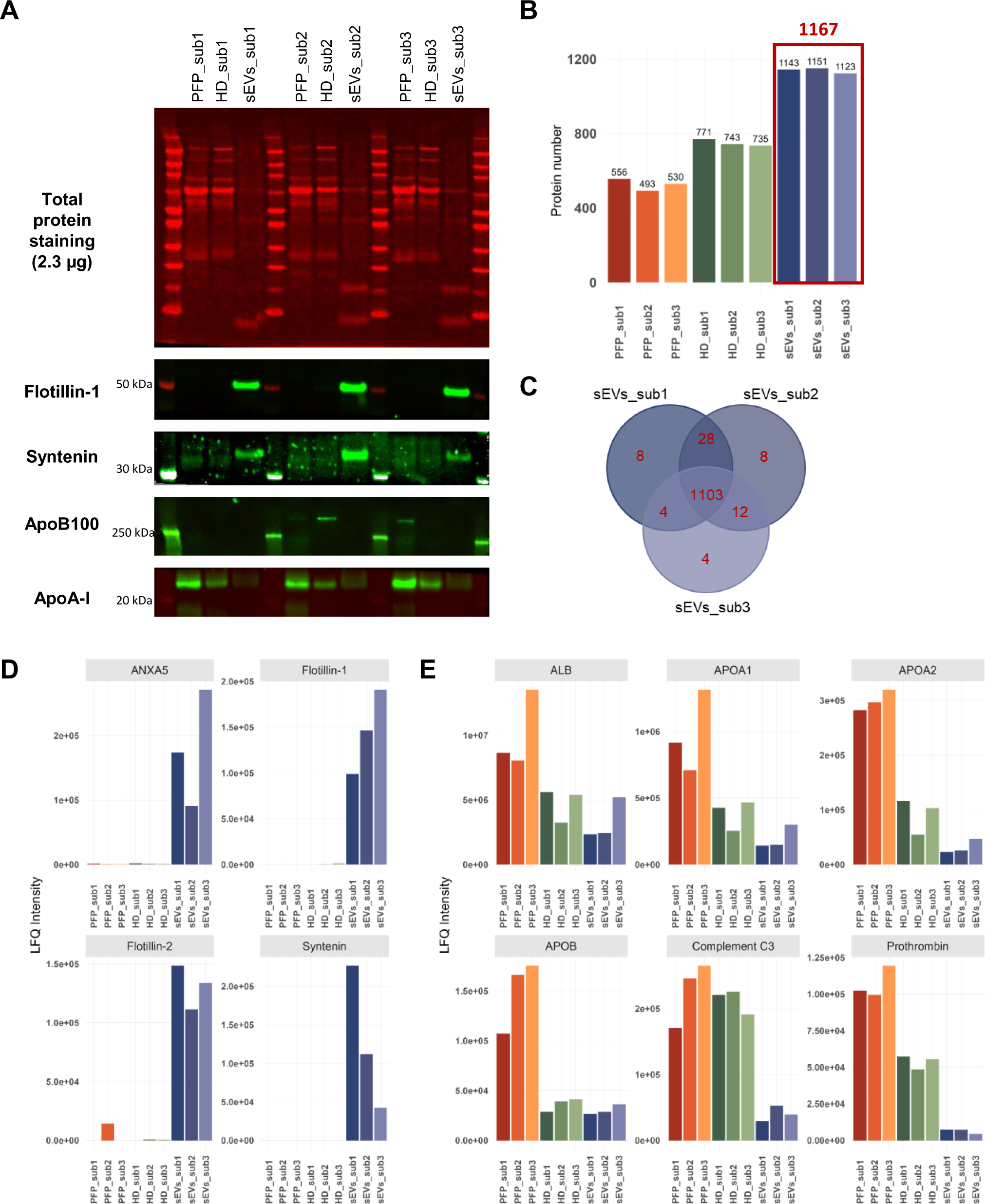
Immunoblot and proteomics characterization of highly enriched plasma sEVs from healthy individuals. **A.** Normalized PFP, HD and sEVs samples (2.3 µg) were electrophoresed for 60 minutes at 160 V. Total proteins were stained, followed by immunoblotting with ApoB100 (LDL marker), ApoA-I (HDL marker) and syntenin and flotillin-1 (general EVs markers). **B.** The number of proteins identified in each sample (PFP, HD and sEVs). The red frame and number indicate total proteins identified in the sEVs. **C.** Venn diagram of identified proteins in plasma sEVs from the three individuals. **D.** The LFQ comparison of representative EVs markers. **E.** The LFQ comparison of plasma proteins and lipoproteins. PFP = platelet free plasma, HD = high density band, sEVs = small extracellular vesicles, sub=volunteer subject.

Gene ontology enrichment analysis of sEVs further validated the identity of the sEVs (Figure 7A), and highlighted their proteomes associated with CNS disease pathways (Figure 7B) the immune system and platelet activity (Figure 7C). We identified neurodegeneration/CNS associated proteins (Figure 7E) such as MBP, SYN1, SYT1, NRGN, TMEM63A, SNCA, SOD1 and NCSTN (full list in Supplemental Table 5) in sEVs. Moreover, proteins associated with mitochondria (82 proteins), the endosomal-autophagic-lysosomal (EAL) pathways (107 proteins) in sEVs (Figure 7E and 7F respectively), highlights the interplay of these intracellular pathways with sEV biogenesis. 26 lipid metabolism proteins were also identified in sEVs (Figure 7G), indicating a potential role for sEVs in regulating lipid homeostasis in recipient cells. Notably, these proteins were either undetectable or present at substantially low levels in PFP. The sEV proteins associated with these pathways are provided in Supplemental Table 5. When comparing the sEV proteome with the Human Plasma Proteome [39], 105 proteins were detected in our sEVs that have not been reported in the Human Plasma Proteome (Figure 7H and Supplemental Table 6). These were predominantly CNS, mitochondrial, histone and EAL associated proteins. In summary, the removal of blood contaminants and lipoprotein particles reduced the complexity of the sample and increased the identification of proteins associated with sEVs, especially those in low abundance such as mitochondria, endosome, lysosome and autophagy associated proteins.

**Figure 7.**
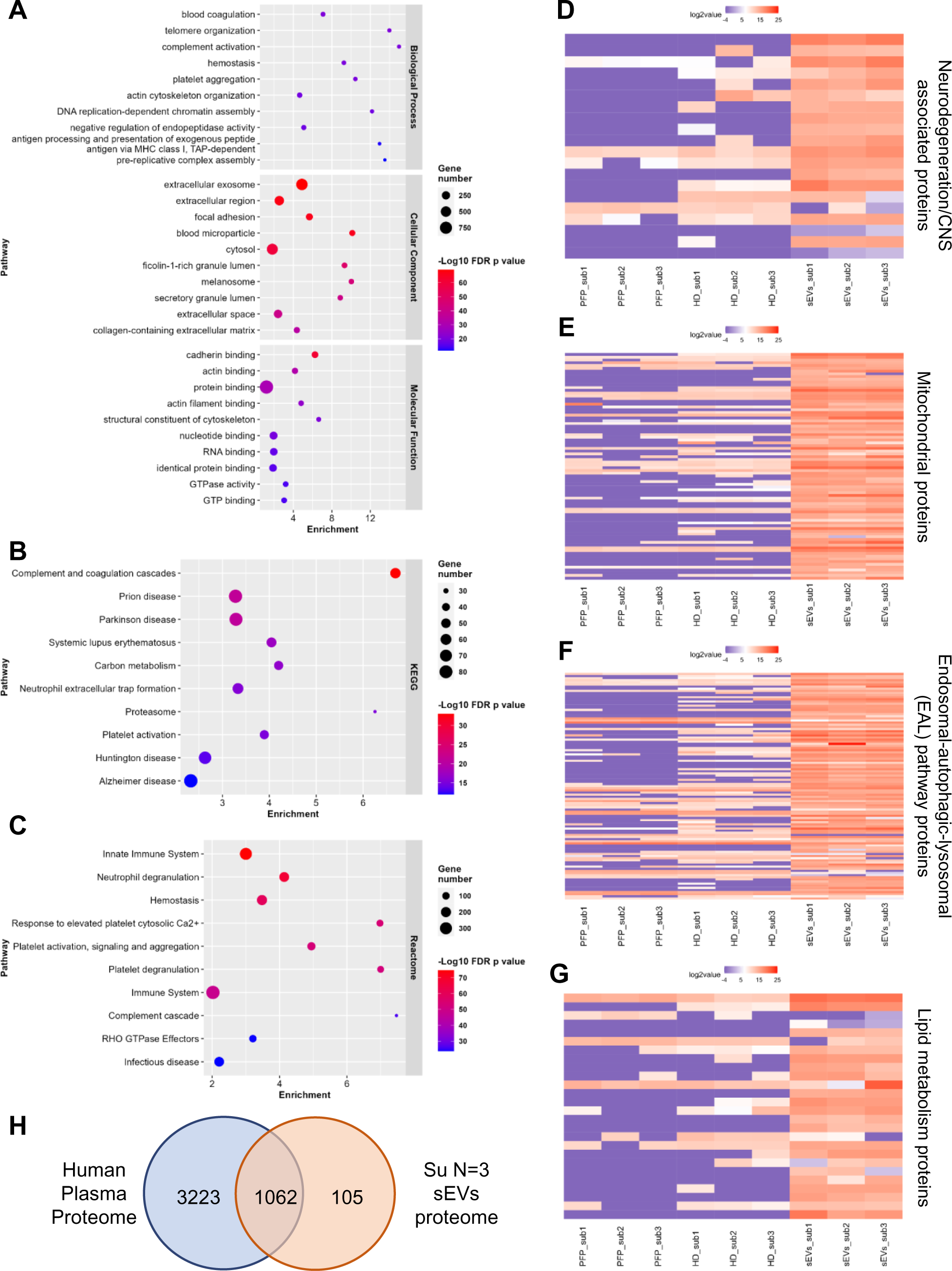
Enrichment analysis of the proteome of highly enriched plasma sEVs from 3 individuals. **A.** Gene ontology enrichment analyses including Top 10 biological process, cellular component and molecular function pathways. **B.** Top 10 enriched KEGG pathways. **C.** Top 10 enriched Reactome pathways. **D.** Venn diagram showing numbers of proteins identified in plasma sEVs compared with Human Plasma Proteome (Deutsch *et al.*, 2021). Histograms show identification and enrichment of **(E)** neurodegeneration/CNS associated proteins, **(F)** mitochondrial proteins, **(G)** endosomal-autophagic-lysosomal (EAL) pathway proteins and **(H)** lipid metabolism proteins in sEVs. The databases were from Human Protein Atlas databases and KEGG pathways. PFP = platelet free plasma, HD = high density band, sEVs = small extracellular vesicles, sub=volunteer subject.

### Lipid profiling of sEVs isolated from three young and healthy female volunteers using DGUC and qEVoriginal™ 70nm SEC column

Given the enrichment of lipids in sEVs and their biomarker and therapeutic potential in diseases, we also examined the lipid content of the highly enriched combined plasma sEV fractions from the three individuals (Figure 8). A total of 612 lipid ions were identified (summarized in Supplemental Table 7). The number of lipids and the mean summed lipid semi-quantitative abundances of lipid categories and lipid classes, expressed in mol% total lipid and mol% lipid class, are summarized in Supplemental Table 8. Plasma sEVs were significantly enriched in SP lipids (40.08 mol% total lipids) and depleted in CE lipids (2.19 mol% total lipids) (Summarized in Figure 8A). For the GP lipid category, sEVs exhibited a significant increase in PE (9.91 mol% total lipids) and PS (14.50 mol% total lipids) lipids and a corresponding decrease in PC lipids (Figure 8B). The observed enrichment in SP lipids was notably contributed by a significant increase in SM lipids, which made up 33.66 mol% total lipids, becoming the most abundant lipid class in sEVs (Figure 8C). Finally TG lipids were depleted in the EVs compared to PFP (Figure 8D). The observed decrease in total CE lipids (16 fold reduction in comparison to PFP) and the increase in SP lipids in sEVs were further illustrated through the depletion of individual CE lipids, CE(18:1), CE(18:2), CE(22:6), CE(14:0) and the enrichment of individual SP lipids, SM(d42:2), SM(d42:1), SM(d34:1), SM(d40:1), Cer(d42:0), and Cer(d35:3), as shown in Figure 8E. In summary, by enriching plasma sEVs via the removal of blood proteins and lipoprotein particles, the bona fide protein and lipid compositions of plasma sEVs can be revealed.

**Figure 8.**
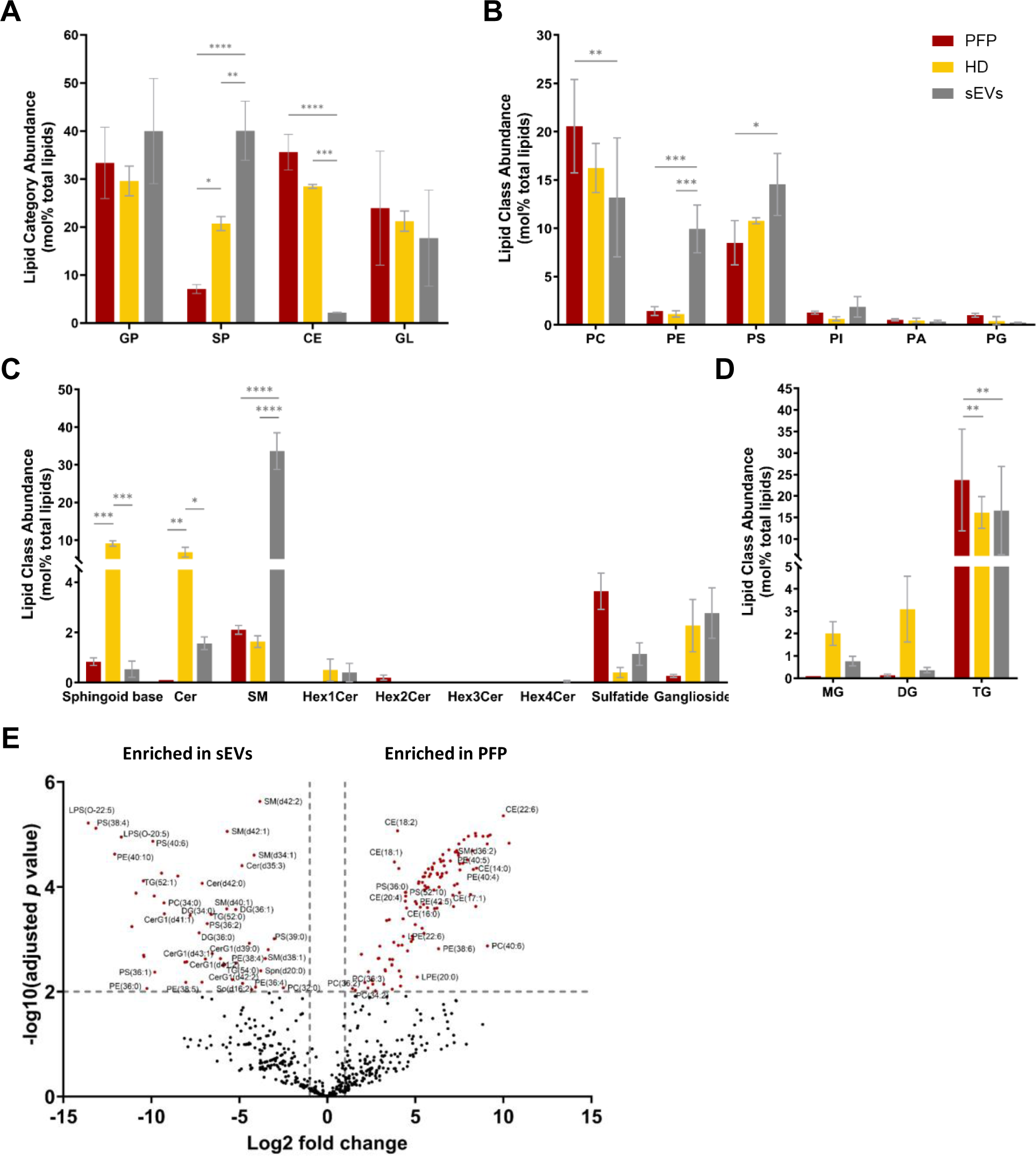
Comparison of lipid abundances between PFP, HD, and sEVs from 3 individuals. **A.** Mol% total lipid abundance distributions at the lipid category level. Four lipid categories, covering glycerophospholipids (GP), sphingolipids (SP), glycerolipids (GL) and sterol lipids, represented by CE. sEVs contained significantly higher levels of SP, and significantly low in CE compared to PFP and HD. **B.** Mol% total lipid abundance distributions at the GP lipid class level. PC, PE, PS, PI, PA and PG. sEVs were lower in PC while enriched in PE and PS compared to PFP. **C.** Mol% total lipid abundance distributions at the SP lipid class level. Sphingoid base, Cer, SM, Hex1Cer, Hex2Cer, Hex3Cer, Hex4Cer, sulfatide and ganglioside. HD was enriched in sphingoid base and Cer while sEVs were specifically enriched in SM when compared to PFP. **D.** Mol% total lipid abundance distribution at the GL lipid class level. MG, DG, TG are included here. Data represent mol% total lipid abundance mean ± standard deviation. Statistical significance was determined by ANOVA followed by Sidak’s multiple comparison test, with multiplicity adjusted *p* value < 0.05, *p < 0.05; **p < 0.01, ***p < 0.001, ****p < 0.0001. **E.** Volcano plot showing comparison between PFP and sEVs at lipid sum composition level (mol% total lipid). Multiple *t* test followed by Benjamin-Hochberg FDR correction was used to determine statistical significance. PFP = platelet free plasma, HD = high density band, sEVs = small extracellular vesicles.

## Discussion

The composition of sEVs in plasma has broad implications for disease diagnosis, and for understanding their associated biological functions, and therapeutic potential. However, the complexity of plasma poses challenges in determining the molecular compositions of sEVs amongst the various other cell-free components that often co-isolate. Here we aimed to provide the research community with a blueprint detailing both the proteome and lipidome of highly enriched sEVs from plasma. We optimized a DGUC with SEC isolation method to dramatically reduce abundant plasma proteins and lipoproteins in sEV preparations. We then applied this approach to plasma obtained from multiple individuals showing enhanced detection of EV proteins associated with mitochondria and EAL pathways and the CNS. Our findings confirm that sphingomyelin (SM) lipids are the predominant lipid species in plasma-derived sEVs. Additionally, we highlight the potential of reporting on the CE content of sEVs, particularly for assessing the effectiveness of isolation methods in eliminating lipoproteins from plasma.

Multiple studies have previously reported the proteomes [27, 36, 40–47] and lipidomes of blood EV preparations [48–50] However, the large majority of isolation methods used in these studies would have co-isolated contaminates. It is well known that separation of abundant plasma proteins and lipoproteins from EVs is not fully achieved by common EV isolation protocols or commercial kits [51, 52]. For molecular characterization therefore, albeit not suitable for use in high-throughput or clinical applications, combining DGUC with SEC is considered one of the most effective methods for obtaining highly enriched EVs [27, 43, 53].

We assessed the purity of the isolated plasma sEVs by monitoring the depletion of plasma proteins such as albumin, and apolipoproteins. Our plasma sEV preparations had minute abundances of albumin (0.9941% total protein abundance) and apolipoprotein (ApoA-I, ApoA-II and ApoB100 combined, 0.0776% total protein abundance). These values are dramatically lower compared to what was reported in a recent meta-analysis investigating the protein profile of plasma EVs (note that our study used data independent acquisition (DIA) where the meta-analysis was from data acquired using data dependent acquisition MS methods) where the albumin level ranged from approximately 10-40% of the total protein abundance in the eight studies included in the meta analysis [51].

We demonstrate that plasma sEVs contain protein networks associated with the intracellular organelles mitochondria, endosome, autophagosome, and lysosome. Some of the proteins have not been detected previously in plasma, or in peripheral sEVs. A number of mitochondria proteins (82 out of 1167 proteins) were enriched in plasma sEVs compared to PFP. Similarly, plasma sEVs were enriched in proteins from EAL pathways (107 out of 1167 proteins). This is presumably due to the crosstalk between sEVs biogenesis and mitochondrial and EAL pathways, which contribute to the cargo loaded into sEVs [5, 6, 54, 55].

While the method we have used does not specifically separate EVs by cell type of origin, or subpopulation, the proteomic data does provide a glimpse into some of the subtypes of EVs in the population. CNS and neurodegeneration associated proteins (MBP, SYN1, SYT1, NRGN, TMEM63A, SNCA, SOD1 and NCSTN for example) were detected, while 46 heart specific proteins, 13 kidney specific proteins, 163 liver specific proteins and 11 lung specific proteins were identified and found to be enriched in plasma sEVs. These proteins were not detected in the platelet-free plasma (PFP) sample, underscoring the advantage of enriching sEVs to high purity from peripheral sources. This enrichment enhances the capability of detecting less abundant proteins through mass spectrometry-based proteomics. As a fundamental element of sEVs, lipids have gained increasing interest in terms of understanding their biological roles, biomarker potential and therapeutic value. Previous studies reporting the lipid content of plasma or sera EV preparations have commonly employed non-ideal isolation methodologies [56] which do not deplete major blood non-EV lipids. The presence of these non-EV lipids can overshadow the bona fide lipids that constitute sEVs, posing a challenge for the detection of low abundant sEV lipids with biomarker potential through MS. In previously published blood EV lipid studies, CE lipids have either not been identified, or reported [49, 50]. Considering that lipoproteins comprise an inner core of CE, the CE lipid content in blood sEV preparations can serve as an indicator of lipoprotein content. Our study has reported on the CE content of isolated sEVs, demonstrating a substantial 16-fold decrease in CE lipids in sEVs compared to platelet free plasma. This finding suggests that we have effectively depleted lipoproteins from blood upon EV isolation. We encourage future studies in blood EV lipidomics to consider reporting the CE content of their sEV preparations, particularly when aiming to assess the efficacy of their isolation methodology in removing lipoproteins. We acknowledge that lipoprotein particles can interact with sEVs and form a corona on the surface of EVs [57, 58] H, hwever our study suggests that the majority of lipoproteins in plasma are non-sEV associated.

EVs isolated from sources other than blood, such as cell lines, urine and adipose tissue, are known to be rich in SM, PS and PE lipids [5, 29, 59–65]. Here, we show that SM lipids are the most abundant lipids in plasma sEVs (33.66 mol% total lipids), followed by PS, PC and PE lipids. This conflicts with Sun *et al*, who previously reported that PC are the dominant lipids (nearly 60 mol% total lipids), followed by SM (over 20 mol% total lipids). However, this discrepancy can likely be attributed to potential co-isolation of other plasma components with the EVs due to the isolation technique that was used in that prior study [50]. Another study, conducted by Peterka *et al.* used various mass spectrometric techniques to examine plasma EVs isolated by polymer precipitation [49]. They observed no difference in the SM lipid composition between plasma and EVs. Notably, they also found that TG lipids made up nearly 50% of total lipids in EVs, higher than that in plasma (30% of total lipids). The TG content may not accurately reflect the composition of EVs in plasma but rather suggests contamination typically associated with polymer precipitation EV isolation [49]. SM, PS and ceramide lipids have been flagged as potential biomarkers for numerous diseases, particularly in CNS disorders [66]. For example, SM and ceramide lipids can be early predictors of cognitive decline [67, 68] and ceramide lipids related to apoptosis have implications in Lewy body disorders [69] and cardiovascular disease [70]. Enriching these lipids from plasma, by isolating highly enriched EVs, enhances their signal to noise ratio, thereby possibly improving the sensitivity of detection of these potential biomarkers in clinical applications. High-throughput technologies for EV isolation however will be required for this to become a reality.

This study underscores the necessity of enriching sEVs to high purity from peripheral sources in order to determine their protein and lipid compositions through mass spectrometry-based proteomics. The dataset resulting from this study acts as an essential foundational resource of sEV protein and lipid information, empowering researchers to pinpoint molecules relevant to their interests for biomarker discovery, high-throughput method design, or the development of techniques for isolating tissue/cell type-specific sEVs from peripheral sources.

## Acknowledgement

We thank the phlebotomists from the Florey Institute. We thank the staff and infrastructure within the Mass Spectrometry and Proteomics Facility (MSPF) located in the Bio21 Institute at the University of Melbourne where the proteomic and lipidomic analysis was performed.

## Data Availability

The datasets generated during and/or analysed during the current study are available in ProteomeXchange.

## Ethics Approval

This study was performed in accordance with the National Health and Medical Research Council guidelines. Approval was granted by the Ethics Committee of Melbourne University Application Number 24582.

## Consent to participate

Informed consent was obtained from all individual participants included in the study.

## Supplementary Figure Legends

**Supplementary Figure 1. Reproducibility of S-Trap proteomics sample preparation and DIA-proteomic analysis method.** The samples, PFP, HD, 6B_F10, 35nm_F10, 4B_F10 and 70nm_F11 were processed and analyzed in triplicates. A. Boxplot shows the Log2 MS2 quantity distribution of the measured samples. B. The number of protein identifications from measured samples in triplicate.

**Supplementary Figure 2. Comparison of the DGUC SEC isolated plasma sEVs proteome with the EV proteomes reported by others.** A. Venn diagram showing numbers of sEVs proteins identified in this study (using DGUC and the 70nm SEC) compared with proteins identified in EVs by Karimi *et al.*, 2018 and Muraoka *et al.*, 2022. B. The comparison of top 10 cellular component enrichment pathways of the proteins associated with the sEVs in this study with Karimi *et al.*, 2018 and Muraoka *et al.*, 2022. C. The interactive networks of the 177 unique proteins identified in the sEVs plasma proteome. The 3 clusters were formed after performing clustering with the KMEANS method.

**Supplementary Figure 3. Lipid metabolism proteins and pathways identified in PFP, HD and EV enriched fractions from density gradient ultracentrifugation (DGUC) followed by size exclusion chromatography (SEC).** Collectively, a total of 65 lipid metabolism proteins were identified from all samples and all were present in the qEVoriginal™ 70nm EV enriched fractions F7-9. 6B EV = Sepharose™ CL-6B fractions F8 and F9; 35nm EV = qEVoriginal™ 35nm fractions F7-9; 4B EV = Sepharose™ CL-4B fractions F8 and F9; 70nm EV= qEVoriginal™ 70nm fractions F7-9.

**Supplementary Figure 4. Identification and enrichment of (A) neurodegeneration/CNS, (B) mitochondria associated proteins, (C) endosomal-autophagic-lysosomal (EAL) pathway, and (D) lipid metabolism proteins in PFP, LD, HD and particle fractions collected from density gradient ultracentrifugation (DGUC) followed by size exclusion chromatography (SEC).** Low abundant EV-associated proteins are minimally detectable via MS of PFP, LD or HD were identified, and some were found enriched in sEVs enriched fractions, specifically sEVs isolated using the qEVoriginal™ 35nm and 70nm SEC columns. The databases were from Human Protein Atlas databases and KEGG pathways. PFP = platelet free plasma, LD= low density band, HD = high density band, 6B = Sepharose™ CL-6B, 35nm = qEVoriginal™ 35nm, 4B = Sepharose™ CL-4B, 70nm = qEVoriginal™ 70nm.

## Supplementary Tables

**Supplementary Table 1.** The protein list and quantity of proteins from PFP, LD, HD, and individual fractions from Sepharose™ CL-6B, qEVoriginal™ 35nm, Sepharose™ CL-4B and qEVoriginal™ 70nm.

**Supplementary Table 2.** The 177 proteins that were uniquely identified in the N=3 sEVs compared to Karimi et al. and the Muraoka et al.

**Supplemental Table 3.** Lipid identification and normalized lipid abundance of PFP, HD, and qEVoriginal™ 70nm EV enriched fractions F7, F8 and F9.

**Supplemental Table 4.** Comprehensive lipid category and class comparisons and statistical results of the PFP, HD, and qEVoriginal™ 70nm EV enriched fractions F7, F8 and F9.

**Supplemental Table 5.** The protein list of neurodegeneration/CNS associated proteins, mitochondrial proteins, endosomal-autophagic-lysosomal (EAL) pathway proteins and lipid metabolism proteins identified in n=3 EV as shown in histograms in Figure 7.

**Supplemental Table 6.** The unique 105 proteins identified in EV from n=3 subjects compared to Human Plasma Proteome Project database

**Supplemental Table 7.** Lipid identification and normalized lipid abundance of PFP, HD, and EV from n=3 young healthy female volunteers.

**Supplemental Table 8.** Comprehensive Lipid category and class comparisons and statistical results of PFP, HD, and EV from n=3 young healthy female volunteers.

