## Supplementary Figures for "Multi-omics characterization of highly enriched human plasma extracellular vesicles"

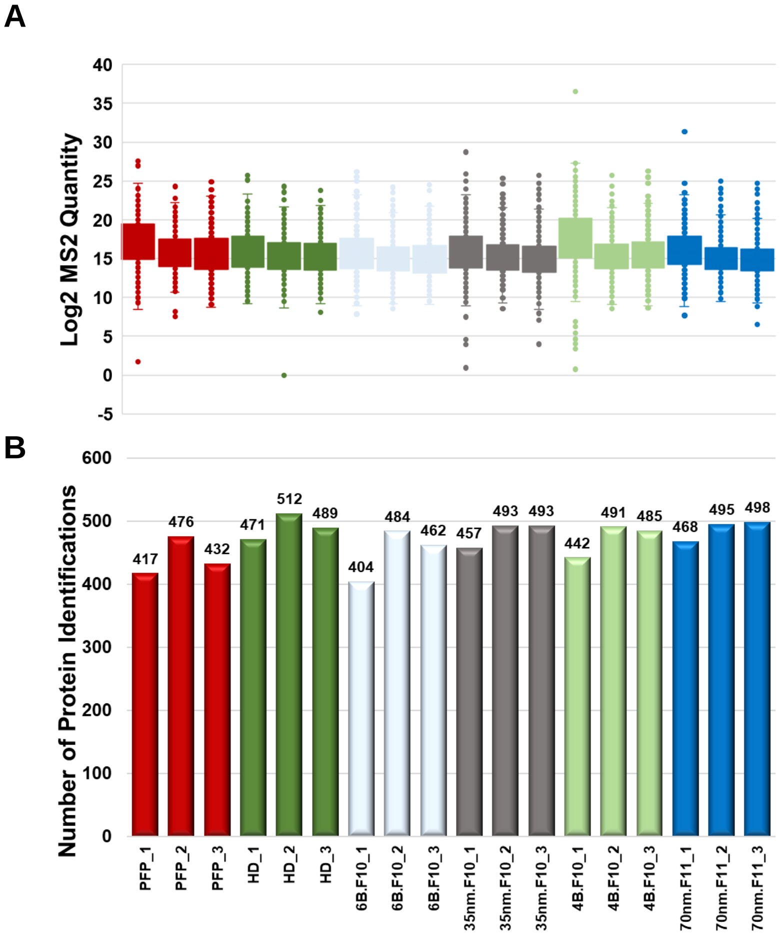


**Supplementary Figure 1. Reproducibility of S-Trap proteomics sample preparation and DIA-proteomic analysis method.** The samples, PFP, HD, 6B_F10, 35nm_F10, 4B_F10 and 70nm_F11 were processed and analyzed in triplicates. **A.** Boxplot shows the Log2 MS2 quantity distribution of the measured samples. **B.** The number of protein identifications from measured samples in triplicate.

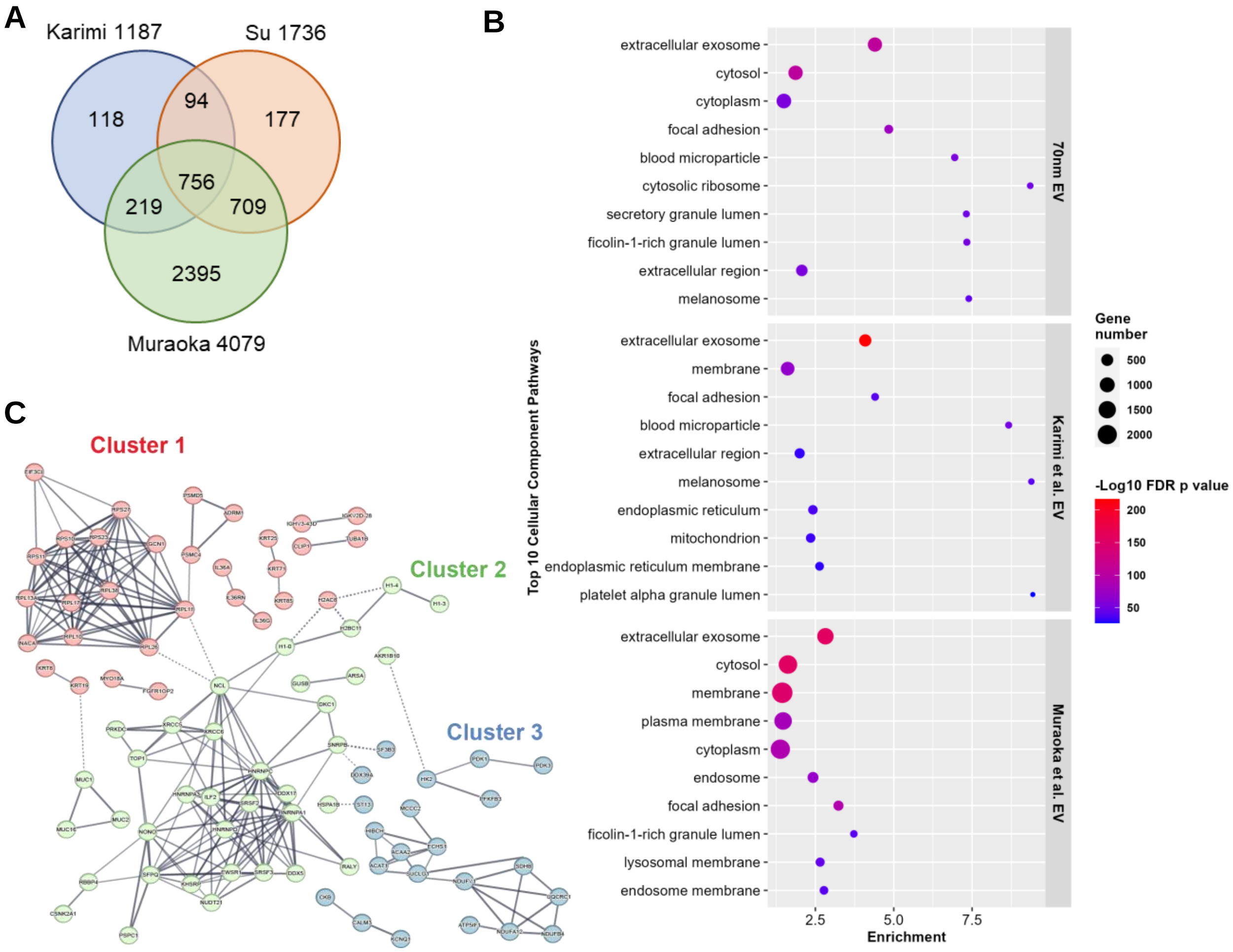


**Supplementary Figure 2. Comparison of the DGUC SEC isolated plasma sEVs proteome with the EV proteomes reported by others. A.** Venn diagram showing numbers of sEVs proteins identified in this study (using DGUC and the 70nm SEC) compared with proteins identified in EVs by Karimi *et al.*, 2018 and Muraoka *et al.*, 2022. **B.** The comparison of top 10 cellular component enrichment pathways of the proteins associated with the sEVs in this study with Karimi *et al.*, 2018 and Muraoka *et al.*, 2022. **C.** The interactive networks of the 177 unique proteins identified in the sEVs plasma proteome. The 3 clusters were formed after performing clustering with the KMEANS method.

| | Number of lipid metabolism proteins |
|---|---|
| PFP | 1 |
| HD | 7 |
| 6B EV | 23 |
| 35nm EV | 63 |
| 4B EV | 45 |
| 70nm EV | **65** |

| Lipid Metabolism Pathway | Number | Lipid Enzymes |
|---|---|---|
| Arachidonic acid metabolism | 13 | ALOX12, GPX1, ALOX12B, CBR1, PLA2G4D, TBXAS1, GPX3, LTA4H, CBR3, GGT1, PLA2G4A, PTGES, PTGS1 |
| Sphingolipid metabolism | 5 | NEU2, GLB1, ASAH1, ARSA, SPTLC2 |
| Glycerolipid metabolism | 9 | ALDH9A1, LPL, AKR1A1, AKR1B10, ALDH2, GK, MGLL, ALDH7A1, ABHD16A |
| Glycerophospholipid metabolism | 6 | LYPLA1, PLA2G4D, ACHE, PLA2G4A, CDS2, GPD2 |
| Biosynthesis of unsaturated fatty acid metabolism | 8 | HACD4, HACD3, HSD17B4, ACOT7, SCP2, TECR, HSD17B12, ACOX1 |
| Ether lipid metabolism | 5 | PAFAH1B1, PLA2G4D, GDPD3, PLA2G4A, AGPS |
| Fatty acid biosynthesis | 4 | ACSL4, ACSL3, FASN, ACSL5 |
| Fatty acid elongation | 10 | HACD4, HACD3, HADHB, HADHA, ACAA2, ECHS1, HADH, ACOT7, TECR, HSD17B12 |
| Fatty acid degradation | 19 | ACSL4, ACSL3, ALDH2, ACADM, ADHX, ACAD, THIL, ECHM, ECHA, THIM, ACDSB, AL9A1, AL7A1, ACADV, CPT1, ECHB, ACOX1, HCDH, ACSL5 |
| Steroid biosynthesis / Steroid hormorne biosynthesis | 2 | LSS, EBP |
| Alpha linolenic acid metabolism | 3 | PLA2G4A, ACOX1, PLA2G4D |
| Linoleic acid metabolism | 2 | PLA2G4A, PLA2G4D |
| Primary bile acid synthesis | 2 | ST2B1, COMT, DHB12, DHB11 |

**Supplemental Figure 3. Lipid metabolism proteins and pathways identified in PFP, HD and EV enriched fractions from density gradient ultracentrifugation (DGUC) followed by size exclusion chromatography (SEC).** Collectively, a total of 65 lipid metabolism proteins were identified from all samples and all were present in the qEVoriginal™ 70nm EV enriched fractions F7-9. 6B EV = Sepharose™ CL-6B fractions F8 and F9; 35nm EV = qEVoriginal™ 35nm fractions F7-9; 4B EV = Sepharose™ CL-4B fractions F8 and F9; 70nm EV= qEVoriginal™ 70nm fractions F7-9.

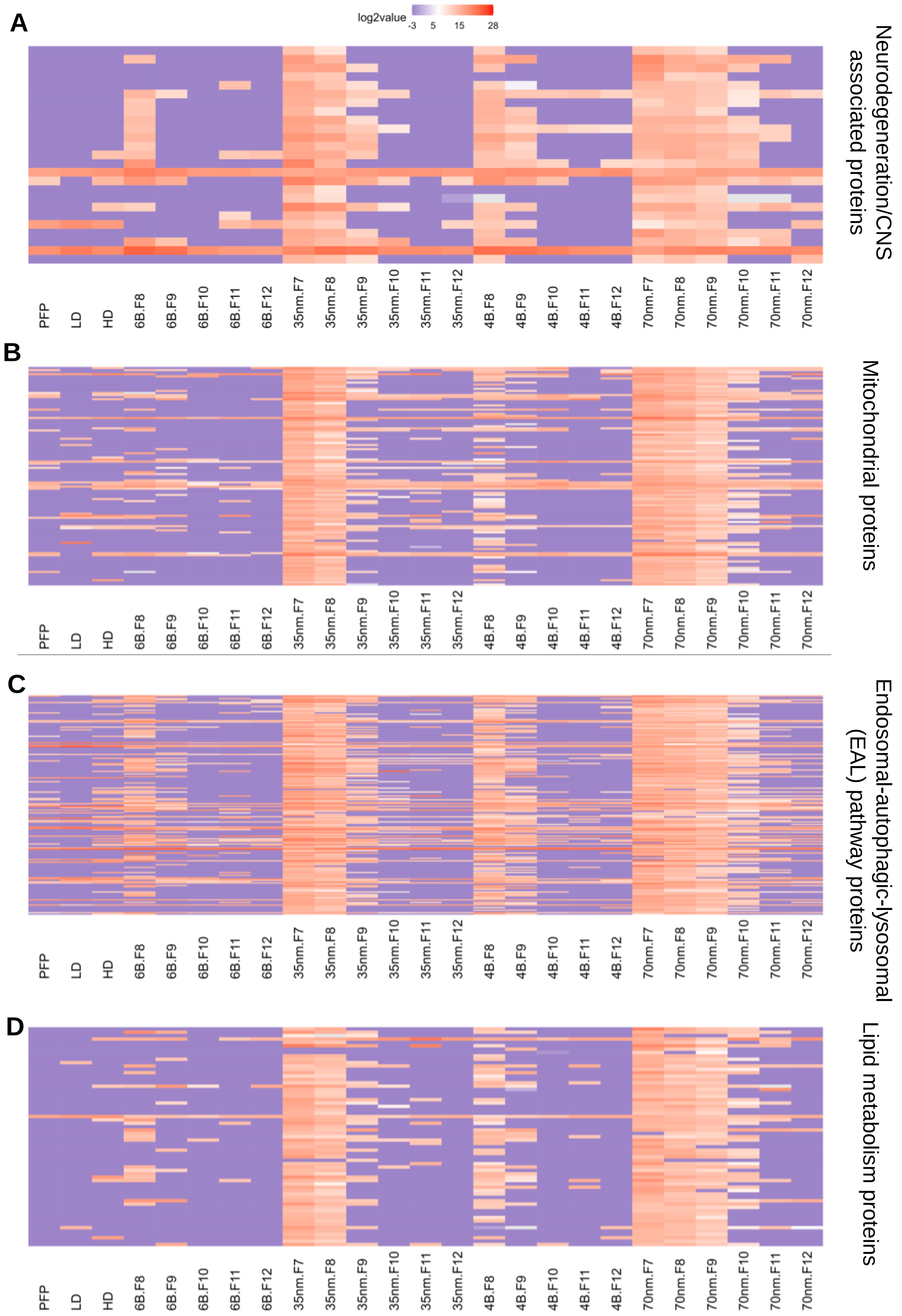


**Supplemental Figure 4. Identification and enrichment of (A) neurodegeneration/CNS, (B) mitochondria associated proteins, (C) endosomal-autophagic-lysosomal (EAL) pathway, and (D) lipid metabolism proteins in PFP, LD, HD and particle fractions collected from density gradient ultracentrifugation (DGUC) followed by size exclusion chromatography (SEC).** Low abundant EV-associated proteins are minimally detectable via MS of PFP, LD or HD were identified, and some were found enriched in sEVs enriched fractions, specifically sEVs isolated using the qEVoriginal™ 35nm and 70nm SEC columns. The databases were from Human Protein Atlas databases and KEGG pathways. PFP = platelet free plasma, LD= low density band, HD = high density band, 6B = Sepharose™ CL-6B, 35nm = qEVoriginal™ 35nm, 4B = Sepharose™ CL-4B, 70nm = qEVoriginal™ 70nm.
